# Differentiation independent neuroprotective role of Vcan^+^ oligodendrocyte precursor cells in poststroke cognitive impairment recovery

**DOI:** 10.1101/2020.11.07.356311

**Authors:** Shanghang Shen, Yongjie Wang, Ruixue Xia, Fanning Zeng, Jun Cao, Xiaolei Liu, Yunbin Zhang, Zhanxiang Wang, Tao Tao

## Abstract

The cell type-specific molecular pathology of post-stroke cognitive impairment (PSCI) in the hippocampus has not been thoroughly elucidated. We analyzed 27,069 cells by using single-cell RNA sequencing, and four oligodendrocyte precursor cell (OPC) subtypes were identified, Vcan^+^ OPCs, which were determined to be the primary cluster among them. Additionally, we examined the features of endothelial cells (ECs) and found that Lcn2^+^ ECs might play neuroprotective roles via Vwf after stroke. These results may facilitate further studies attempting to identify new avenues of research and novel targets for PSCI treatment.

---

Poststroke cognitive impairment (PSCI) is a major sequela in cerebral ischemia (CI) patients. The prevalence of PSCI ranges from 20% to 80%, and varies depending on country, race, and diagnostic criteria^1^. PSCI can result in worse life quality, lower survival rate, and heavier social burden. The management and mechanisms of PSCI have recently attracted increasingly concentration ^2–5^. Hippocampal lesions have been proposed to play a pivotal role in PSCI^1^. However, the cell type-specific molecular pathology of PSCI has not been elucidated.

In this study, we adopt a mouse two-vessel occlusion (2VO) ischemia model to mimic CI-induced cognitive impairment, which presented significant cognitive decline 7 days after surgery regarding both escape latency and pathlength, but not speed in the training stage of the Morris water maze test (Fig. 1B-D). Moreover, the time spent in the target quadrant of the 2VO group was decreased compared with that of the sham group (Fig. 1E), indicating impairment of the hippocampus. After filtering, 27,069 single cell gene expression profiles (12108 from the sham group and 14961 the from PSCI group) were projected onto two dimensions via uniform manifold approximation and projection (UMAP)^6^. Thirty-two clusters were obtained, and no significant bias between the two groups was observed (Fig. 1F, Table S1). Based on known cell marker expression, 13 cell types of neuronal, glial and vascular lineages were identified (Fig. 1G, sFig. 1, sFig. 2). The proportions of 13 cell types were compared between the two groups, and vascular endothelial cells and oligodendrocytes accounted for the majority of the cells. As expected, astrocytes, neurons and oligodendrocytes showed the most striking decrease, while vascular endothelial and smooth muscle cells were increased in PSCI (Fig. 1H). Cell numbers and genes detected in each cell type are shown in Fig. 1I. We also identified novel markers for each hippocampal cell type (Fig. 1J, Table S2). Next, we investigated cell type specific molecular changes by performing differential expression gene (DEG) analysis. Interestingly, astrocytes (ASTs), ependymal cells (EPNs), glutamatergic neurons (GLUTNs), oligodendrocyte precursor cells (OPCs) and pericytes (PERs) exhibited the highest number of DEGs, while macrophages (MACs) exhibited almost no changes in DEGs (Fig. 1K). We also identified cell type-specific DEGs (Fig. 1L, Table S3), e.g., *Ler2* for ASTs, *Fth1* for ECs, *Cnr1* and *Sox4* for GABANs, and *Reln* for GLUTNs. Additionally, common DEGs in at least 2 cell types (e.g., *Hspa1b, mt-Atp6, Neat1* and *Lars*) were identified (Fig. 1M and Table S4). Finally, GSEA based hierarchical clustering was employed, and 13 cell types showed distinct hallmark patterns (Fig. 1N).

**Figure 1.**
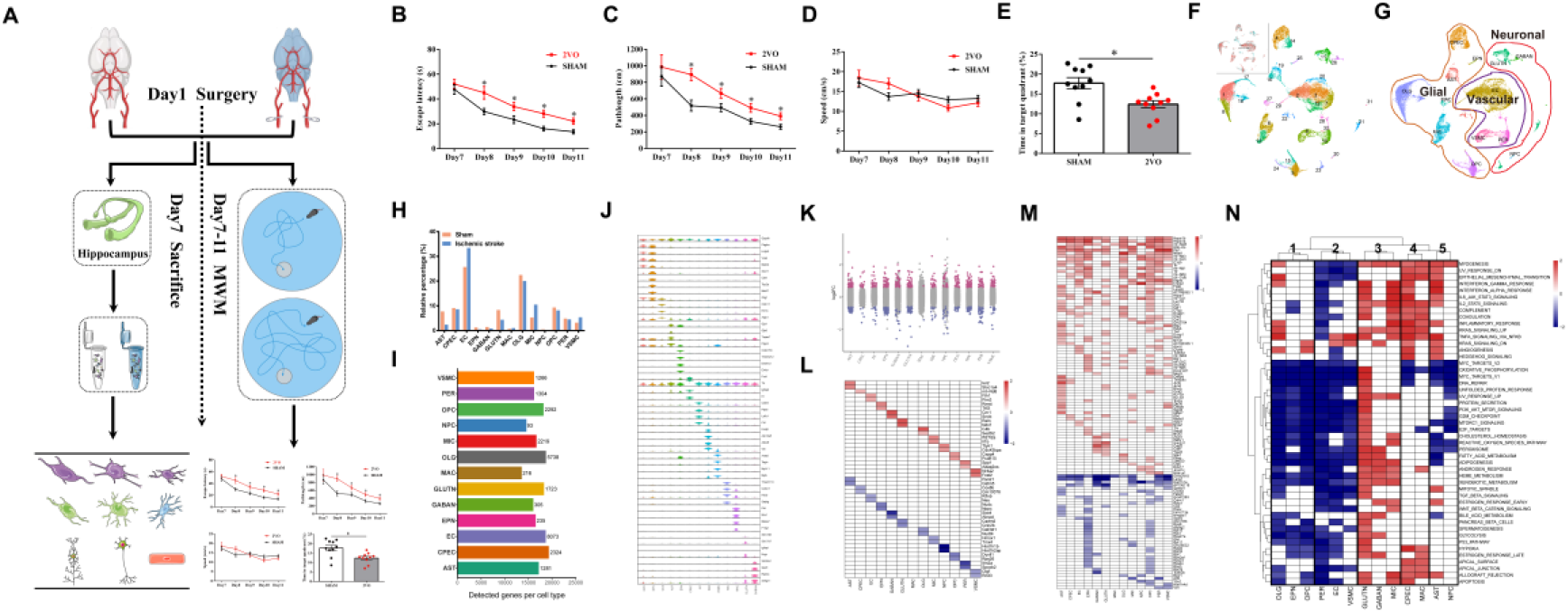
Hippocampal heterogeneity and molecular alterations of 13 cell types in ischemic stroke. (A) Workflow of this study; (B) The Morris water maze test was performed with days indicated. Escape latency to the platform (B) was measured during the acquisition test at 7, 8, 9, 10, and 11 d after 2VO surgery. Pathlength (C) and locomotion speed (D) were also recorded and plotted (Sham: n = 10; 2VO: n = 10). (E) The percentage of time spent in the target quadrant during the spatial exploration stage of the Morris water maze probe test (Sham: n = 10; 2VO: n = 10). Data are presented as the mean ± SEM. Error bars indicate SEM. * indicates a significant difference between the Sham and 2VO groups; *P < 0.05; Test used in B – D RM 2 ANOVA with Bonferroni post hoc test; E Student’s *t*-test. (F) Thirty-two clusters of (G) 13 cell types of neuronal, glial and vascular cells were identified; (H) cell type percentage, (I) numbers and genes in ischemic stroke and sham groups; (J) known and novel markers were identified; (K) overall DEGs and (L) cell type specific DEGs in ischemic stroke; (M) common DEGs in at least 2 cell types; (N) functional alterations of 13 cell types.

Neurogenesis is the process of producing new functional neurons from other cell types, including the proliferation and differentiation of progenitor cells into mature neurons. In cerebral ischemic stroke, enhanced neurogenesis has been reported after stroke^7^, suggesting a potential avenue for ischemic stroke therapy. Therefore, we focused on the cell cycle state of 13 cell types (sFig. 3, Table S5). As expected, 78.72% of cluster 30, NPCs, expressed genes in G2/M phase, and 21.28% expressed genes in S phase, with no cells being in G1 phase (Fig. 2A). Notably, a large proportion of cluster 5 (OPCs) were in S phase. OPCs are multipotent for differentiation into other cell types, including oligodendrocytes and neurons^8^. Consistent with previous reports^9–10^, we identified four OPC subtypes (Fig. 2B) with novel markers for each subtype, such as *Cspg5* for OPC-1, *Vcan* for OPC-2, *Fyn* for OPC-3 and *Pclaf* for OPC-4 (Fig. 2C, Table S6). To further explore cell trajectories within OPC development, Slingshot^11^ pseudotime analysis showed a developmental trajectory from OPC-4 to OPC-1, then to OPC-2, and finally to OPC-3 (Fig. 2D), suggesting OPC-4 (*Pclaf*_+_ OPC) as a primary cluster. As OPC-3 (Fyn^+^ OPCs) expressed high levels of Plp1 and Mbp (Table. S6), important modulator in myelin formation, it was defined as a cluster that committed differentiation into oligodendrocytes. Further analysis integrating OPC-3 and oligodendrocytes also supported this conclusion (Fig. 2E), with the development route of OPC-3/OLG-17/OLG-8/OLG-1/OLG-18. OPC-1 (Cspg5^+^ OPCs) strongly express the immature neuron marker Sox11, and cluster 23 highly expresses the mature neuron marker Syt1. Slingshot analysis revealed a trajectory from OPC-1 (e.g., *Pclaf*, and *Spc24*) to NPCs (e.g., *Birc5*, and *Mki67*) to immature neurons (e.g., *Sox11*) to mature neurons (e.g., *Reln*) (Fig. 2E), implying that Cspg5^+^ OPCs are potential neuron-differentiated OPCs. As many DEGs in OPC-2 (*Vcan^+^* OPCs) are involved in energy metabolism (Fig. S4), which is a well-known role of astrocytes^12^, Slingshot analysis was employed to explore the relationship between OPC-2 and astrocytes. Interestingly, we observed distinct trajectories from *Vcan^+^* OPCs to three astrocyte clusters 22, 19 and 16 (Fig. 2E), suggesting astrocyte commitment of OPC-2 (*Vcan^+^* OPCs). Finally, the proportions of 4 OPC clusters in the sham and PSCI groups were compared, and interestingly, only *Vcan^+^* OPCs exhibited an increase (17.72% to 32.57%) in the PSCI group (Fig. 2F).

**Figure 2.**
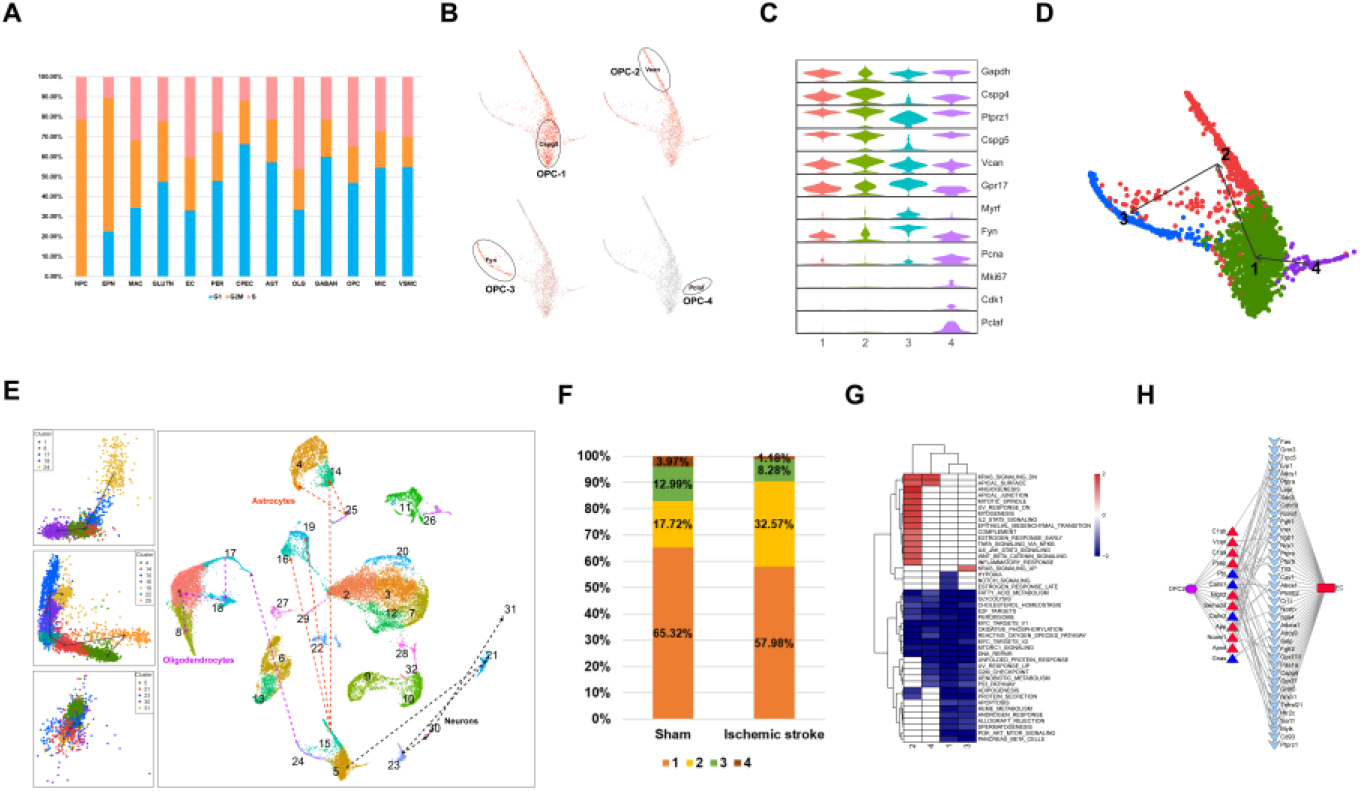
Differentiation route and neuroprotective role of Vcan^+^ OPCs in ischemic stroke. (A) Cell cycle distribution of 13 cell types; (B-C) Known and novel markers of 4 OPC subtypes; (D) Slingshot showing the differentiation route within 4 OPC subtypes; (E) Slingshot reveals the differentiation route of OPCs to oligodendrocytes, astrocytes and neurons; (F) Percentage of 4 OPC subtypes in ischemic stroke and sham groups; (G) GSEA analysis based hierarchical clustering identified OPC2 (Vcan^+^ OPCs) specific pathways; (H-I) Ligand-receptor interaction of Vcan^+^ OPCs with endothelial cells;

Previous studies noted that by secreting pro-angiogenic factors (e.g. VEGF), perivascular OPCs exert an angiogenesis promoting role after brain ischemia^13^. Given that astrocytes were decreased in the PSCI group (Fig. 1D), we hypothesized that OPC2 might function in a differentiation-independent manner. Intriguingly, we noticed that angiogenesis and myogenesis are *Vcan*^+^ OPC-specific processes (Fig. 2G), indeed, vascular endothelial cells and smooth muscle cells are both increased in PSCI (Fig. 1D). We hypothesized that cell-cell communication between *Vcan*^+^ OPCs and vascular cells may play a protective role. Indeed, vascular endothelial cells were found to be important in stroke recovery^14^. In this study, six clusters (i.e., C2, C3, C7, C12, C20 and C29) with higher Cldn5 expression were identified as endothelial cells (ECs) (Fig. 3A, Fig. 1B and Fig. S1). We also identified potential markers for 6 clusters: *Hmcn1* for C2, *Car4* for C3, *ATf3* for C7, *Gkn3* for C12, *Lcn2* for C20 and *Plvap* for C29 (Fig. 3B and C, Table S6). The marker of Lcn2 for brain vascular endothelial cells has been reported in recent studies^15^, and other new endothelial cell markers were proposed in the current study. In keeping with the overall increase in EC after stroke, all 6 EC clusters showed higher levels (Fig. 3D), supporting the neuroprotective role of endothelial cells after stroke. Then, we compared DEGs (Fig. 3E), and employed GSEA functional analysis for 6 EC clusters (Fig. 3F). As C20 showed the most notable increase (1.10% to 1.82%) and DEGs in all 6 EC clusters, we then focused on EC20 (Lcn2^+^ vascular endothelial cells). As shown in Fig. 3F, specifically enriched pathways in EC20 include cholesterol hemostasis, IL6-JAK-STAT3, hypoxia and inflammatory responses. To further investigate the role of EC20 in stroke, we applied ligand-receptor interaction analysis of EC20 with neurons. As shown in Fig. 3G–3I, such ligands as Vwf are proposed to be mediators in EC20 mediated neuroprotection ^16^.

**Figure 3.**
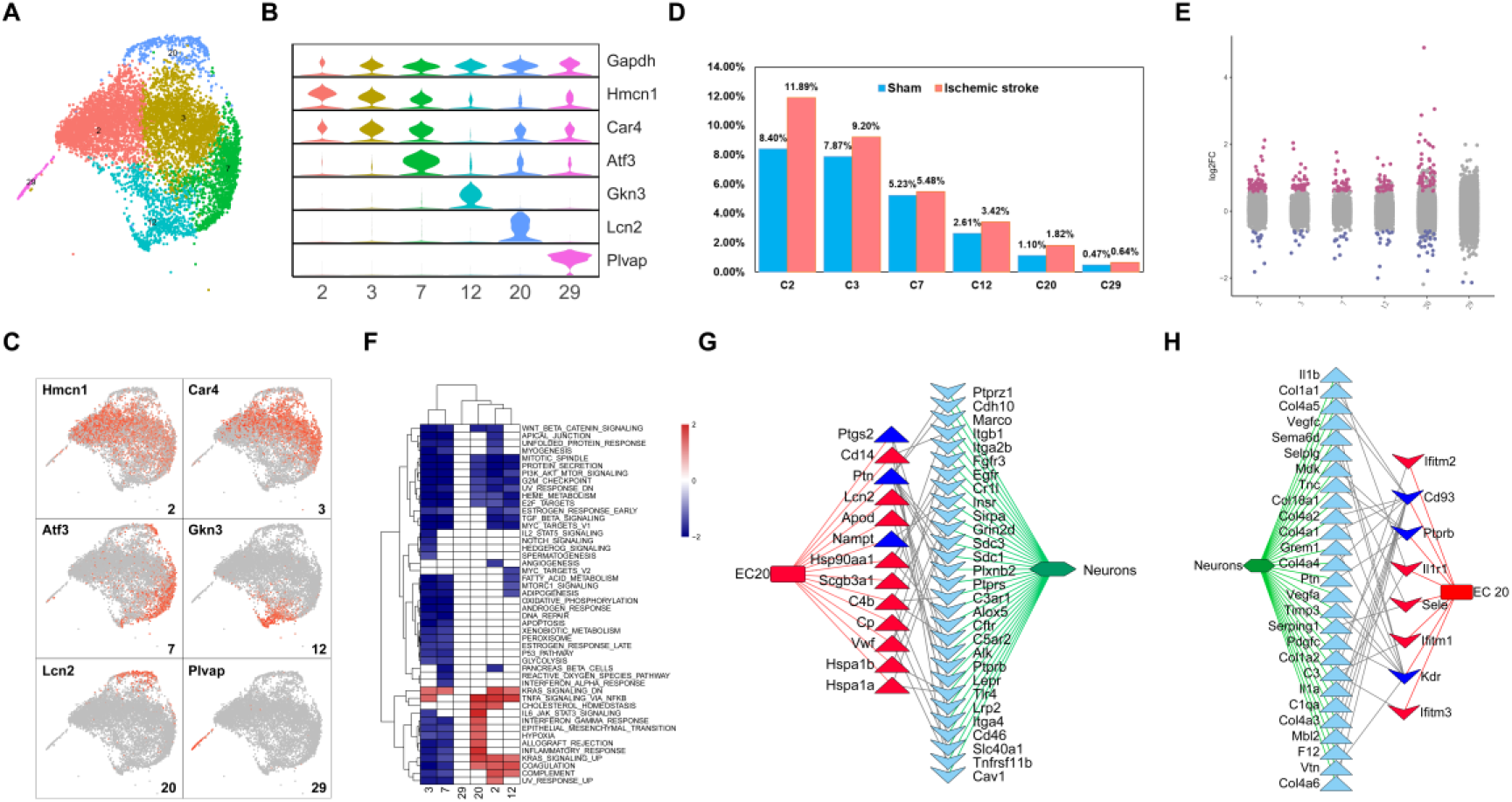
Neuroprotective role of Lcn2^+^ ECs with neurons in ischemic stroke. (A) UMAP showing 6 endothelial cell subtypes; (B-C) Known and novel markers in in 6 EC subtypes; (D) Percentage of 6 EC subtypes in sham and ischemic stroke; (E) DEGs in 6 EC subtypes; (F) GSEA analysis based hierarchical clustering pathways in 6 EC subtypes; (G-H) Ligand-receptor interaction of Lcn2^+^ ECs with neurons;

Taken together, the results of this study indicate neuronal, glial and vascular lineage heterogeneity and cell-specific molecular changes in PSCI. Four subtypes of OPCs with distinct differentiation routes were identified. Notably, Vcan^+^ OPCs may exert a protective role by expressing pro-angiogenesis factors. Additionally, Lcn2^+^ vascular endothelial cells may also protect neurons via Vwf in PSCI. Overall, these results may establish a foundation for further research attempting to identify new avenues of research and novel targets for PSCI treatment.

## Methods

### Mouse two-vessel occlusion (2VO) cerebral ischemia model

All procedures and mouse handling protocol were approved by The Animal Care and Use Committee of Southern Medical University and were in keeping with the guidelines established the International Association for the Study of Pain. Conventional SPF C56BL/7 mice were purchased from Shanghai SLAC Laboratory Animal Center (Shanghai, China). Mice were maintained in a pathogen-free SPFII animal facility in a condition-controlled room (23 ± 1°C, 50 ± 10% humidity). A 12 h light/dark cycle was automatically imposed. Mice were housed in groups of four to five per individually ventilated cage and given access to food and water ad libitum unless otherwise stated in the methods below. researchers were blinded to the animals’ treatments and sample processing throughout the subsequent experimentation and analyses.

Bilateral occlusion of the mouse common carotid arteries was performed as previously reported ^17^. Before surgery, mice were fasted overnight, but were free to drink. The mean arterial blood pressure was measured using a noninvasive blood pressure system (Product #CODA, Kent Scientific Corporation). Mice were anesthetized with isoflurane (5% for induction and 1.5%–2.5% for maintenance). Rectal temperature was maintained at 37.0±0.5 °C during the entire surgical procedure by a rectal temperature probe and a heating pad (Product # TCAT-2 Temperature Controller, Harvard Apparatus, USA). The ventral neck region’s hair was shaved to expose the skin and then disinfected. An incision in the midline of the ventral neck region was made with a scalpel. Then, the superficial fascia was dissected to expose the bilateral common carotid arteries under the operating microscope and both carotid arteries were clamped with microvessel clamps for 50 min to induce global cerebral ischemia. Mouse hippocampal blood flow (hrCBF) was detected by using a laser Doppler flowmetry (moorVMSLDF2, Moor Instruments, Wilmington, DE) with a 0.5 mm flexible fiber optic. The isoflurane level was decreased to 0% 2 min before the release of the microvessel clamps and resumption of blood reperfusion. The neck incision was sutured with 5-0 sterile silk sutures. The animals were moved into a warm (37.0 ± 1 °C) recovery chamber (Product #DW-1, Harvard Apparatus, USA) to prevent postischemic hypothermia. For the sham group mice, microvessel clamping was performed on the bilateral common carotid arteries. The clamping was immediately released to enable instant reperfusion. The subsequent procedure was same as that performed on the cerebral ischemia group, including anesthesia processes.

### Morris water maze

The Morris water maze was employed to determine the spatial learning ability and memory of the mice. The Morris water maze analysis was performed in a white-water pool (120 cm in diameter with 2/3 of transparent water) containing a circular bright black platform (14 cm in diameter and submerged 1.5 cm beneath the water surface). Various patterns were equally distributed around the wall as visual cues for mice during the experiment. The experiment was started on day 7 after 2VO surgery. In the training stage, the spatial acquisition learning ability training consisted of five consecutive days, with every training day comprising four trials with 15 min inter-trial interval. The entry points of mice were randomly selected each time from the different designated locations. Once the mouse successfully found the platform within 60 s, it was placed into a cage under a warming lamp as a reward. Otherwise, the mouse was gently and manually guided to the platform and allowed to remain there for at least 20 s. The escape latency and pathlength to the platform were recorded each day to assess the spatial learning ability.

On day 6 (day 11 after 2VO surgery), a probe trial test was performed. The hidden platform was removed from the pool, and the mouse was placed in the quadrant diagonally opposite the target quadrant and allowed to swim for 60 s freely. The percentage of time spent in the target quadrant was recorded and analyzed as a measure of spatial memory retention. Mice were monitored by a camera, and their trajectory was analyzed using the Smart (V3.0, Panlab Harvard Apparatus).

### Cell isolation

Single cells were obtained according to a previously described procedure with slight modification^18^. Specifically, the individual adult male mouse was deeply anesthetized in an isoflurane chamber and decapitated. The brain was removed, and the whole hippocampus was rapidly dissected. We then wholly cut the tissue thoroughly into pieces and transferred it to a 1.5 mL microcentrifuge tube with 2 mg/mL isolation solution, containing pronase (Sigma, Cat#P6911-1G) and 50 μg/mL DNaseI (Sigma, cat. no. D5025) in 1 mL Hibernate A (Invitrogen, cat. no. A1247501)/B27 (Invitrogen, cat. no. 17504) medium (HABG). The tissue and solution were mixed for 30 min at 37°C in a horizontal shaker at 200 rpm. After incubation, the tissue was gently triturated by polished tips, and single cells were released. The purified cells were obtained by density gradient centrifugation at 800 g for 15 mins, resuspended in 1x PBS (calcium and magnesium-free) containing 1% BSA, and then centrifuged at 200 g for 2 mins. The single cells were concentrated and resuspended in the desired medium. The cell number was counted and viability was measured based on trypan blue staining.

### Single cell RNA sequencing and raw data preprocessing

The cell counts and viability of single-cell suspensions were determined, and samples with cell survival rates above 80% were employed in subsequent procedures. Cells that passed the test were washed and resuspended to prepare a suitable cell concentration of 700~1200 cells/ul for 10x Genomics Chromium ™ system operation. Based on expected number of target cells, GEMs (Gel Bead in Emulsion) were constructed for single cell isolation. After the GEMs were formed, the GEMs were collected and reverse-transcribed in a PCR machine to achieve labeling. Then, GEMs were destroyed, the first strand cDNA was purified and enriched with magnetic beads, and it was subjected to cDNA amplification and quality control. Qualified cDNAs were applied for library construction with 10x Genomics Single Cell 3’ v2 Reagent Kit, and then fragmentation was performed adapters were added, and PCR index of samples was determined. At last, Illumina NovaSeq platform PE150 sequencing mode was conducted for sequencing, and the sequencing volume criteria was set > 50 k reads/cell.

Sequenced samples were processed using the Cell Ranger 3.1.0 pipeline and aligned to the GRCm38 (mm10) mouse reference genome. Cell with fewer than 650 detected genes/cell and genes that were expressed by fewer than 20 cells (0.025% of all cells in the dataset) were removed before identification of variable genes in the dataset, cell centering and scaling.

### Dimensionality reduction and clustering

To visualize and interpret single cell RNA sequencing data, two-dimensional projections of cell populations were achieved by first reducing the dimensionality of the gene expression matrix using principal component analysis (PCA) and, then further reducing the dimensionality of these components using t-distributed stochastic neighbor embedding (t-SNE)^19^and uniform manifold approximation and projection (UMAP)^6^. As tSNE is a nonlinear embedding that does not preserve distances, we used UMAP to analyzed the distances of clusters and cell types in this study.

### Significantly dysregulated genes analysis

To find differentially expressed genes, we used the Mann-Whitney U test, a nonparametric test that detects differences in the level of gene expression between two populations. Using the Mann-Whitney U test, we compared the distributions of expression levels of every gene separately. P values were adjusted using the Bonferroni correction for multiple testing. Genes expression fold changes was calculated as Median (PSCI)/ Median(sham). Significant dysregulated genes were screened using the criteria of absolute value of Foldchange >1.5 and adjusted P value < 0.05.

### GSEA hierarchical clustering analysis of common hallmark pathways and

To identify common hallmark pathways in cell types, OPC and EC clusters, we first applied GSEA enrichment analysis for all genes in corresponding populations. Hallmark gene sets are coherently expressed signatures derived by aggregating many MSigDB (Molecular Signatures Database) gene sets to represent well-defined biological states or processes. After each hallmark pathways was acquired, the Negative normalized Enrichment Score (NES) was applied for hierarchical clustering and heatmap construction for significantly enriched pathways.

### Pseudotime analysis of cell trajectories with Monocle and Slingshot

To investigate cell trajectories of OPCs, we utilized Monocle^20^ and Slingshot^11^ to infer branching lineage assignments and developmental distances. Slingshot is a method for inferring cell lineages and pseudotimes from single-cell gene expression data to identify multiple trajectories. In this study, gene expression data of 4 OPC clusters and clusters in oligodendrocytes, neurons, astrocytes, ependymal cells and CPECs was applied for Slingshot analysis.

### Cell Cycle Phase Assignments

The cell cycle phase analysis was conducted with CellCycleScoring function in Seurat to obtain cell cycle scores and phase assignments. The Cell-Cycle Scoring from Seurat makes available a list of cell cycle phase marker genes for humans and performs phase scoring as described in a the previously published paper^21^.

### Ligand-receptor interaction analysis

The ligand-receptor interaction analysis was primarily based on CellphoneDB, a novel repository of ligands, receptors and their interactions^22^. For OPC2-EC interaction, dysregulated ligands, receptors and interactions with ECs from OPC2 were extracted. For EC20-neurons interaction, dysregulated ligands, receptors and interactions with neurons from EC20 were extracted. Extracted ligands and receptors were presented with Cytoscape v3.6.1.

### Data availability

The scRNA-seq data used in this study would be deposited as the requirements of editorial policy. Raw image files used in the figures that support the findings of this study are available from the corresponding authors upon reasonable request.

### Data analysis

GraphPad Prism 8 was used for data processing and analysis. All data were expressed as means ± SEM. We used unpaired *t*-test, paired *t*-test, one-way analysis of variance (1-way ANOVA), or two-way analysis of variance (2-way ANOVA) to conduct data analysis. All data were presented as the mean ±SEM in all cases, there was a statistical significance in p values less than 0.05.

## Acknowledgments

We are grateful for the help of Litao Li from Heibei general hospital. This study was supported by the National Natural Science Foundation of China (81801102 and 81302758), the Science and Technology Planning Project of Guangdong Province, China (Grant No. 2017A020215133, 2014A020212170), and the Medical Interdisciplinary Research Funding of Henan University (Grant NO. CJ1205A0240011).

## Author contributions

S. Shen and Y. Wang performed single-cell isolation. R. Xia performed behavioral data analysis. F. Zeng, J. Cao and X. Liu carried out behavioral test. Y. Wang and Y.Zhang performed the sc-RNA seq data analysis. Y. Wang, Y.Zhang, Z. Wang and T. Tao designed experiments, performed data analysis, approved the draft, and wrote the paper.

---

**Supplementary Fig. 1.**
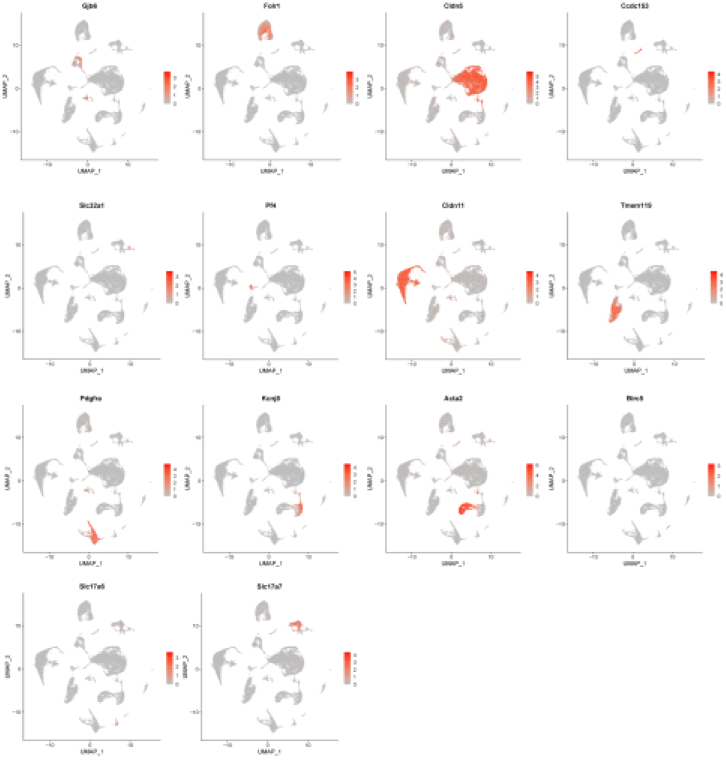
UMAP visualization of each cell type showing the expression of representative well-known marker genes. That is, Gjb6 for astrocytes (ASTs), Folr1 for choroid plexus epithelial cells (CPECs), Cldn5 for endothelial cells (ECs), Ccdc153 for ependymocytes (EPNs), Slc32a1 for GABAergic neurons (GABANs), both Slc17a6 and Slc17a7 for glutamatergic neurons (GLUTNs), Cldn11 for mature oligodendrocytes (OLGs), Pdgfra for oligodendrocyte precursor cells (OPCs), Birc5 for neural progenitor cells (NPCs), Kcnj8 for pericytes (PERs), Acta2 for vascular smooth muscle cells (VSMCs), Tmem119 for microglia (MICs), and Pf4 for macrophages (MACs). Numbers reflect the actual nUMI detected in each cell for the specified gene.

**Supplementary Fig. 2.**
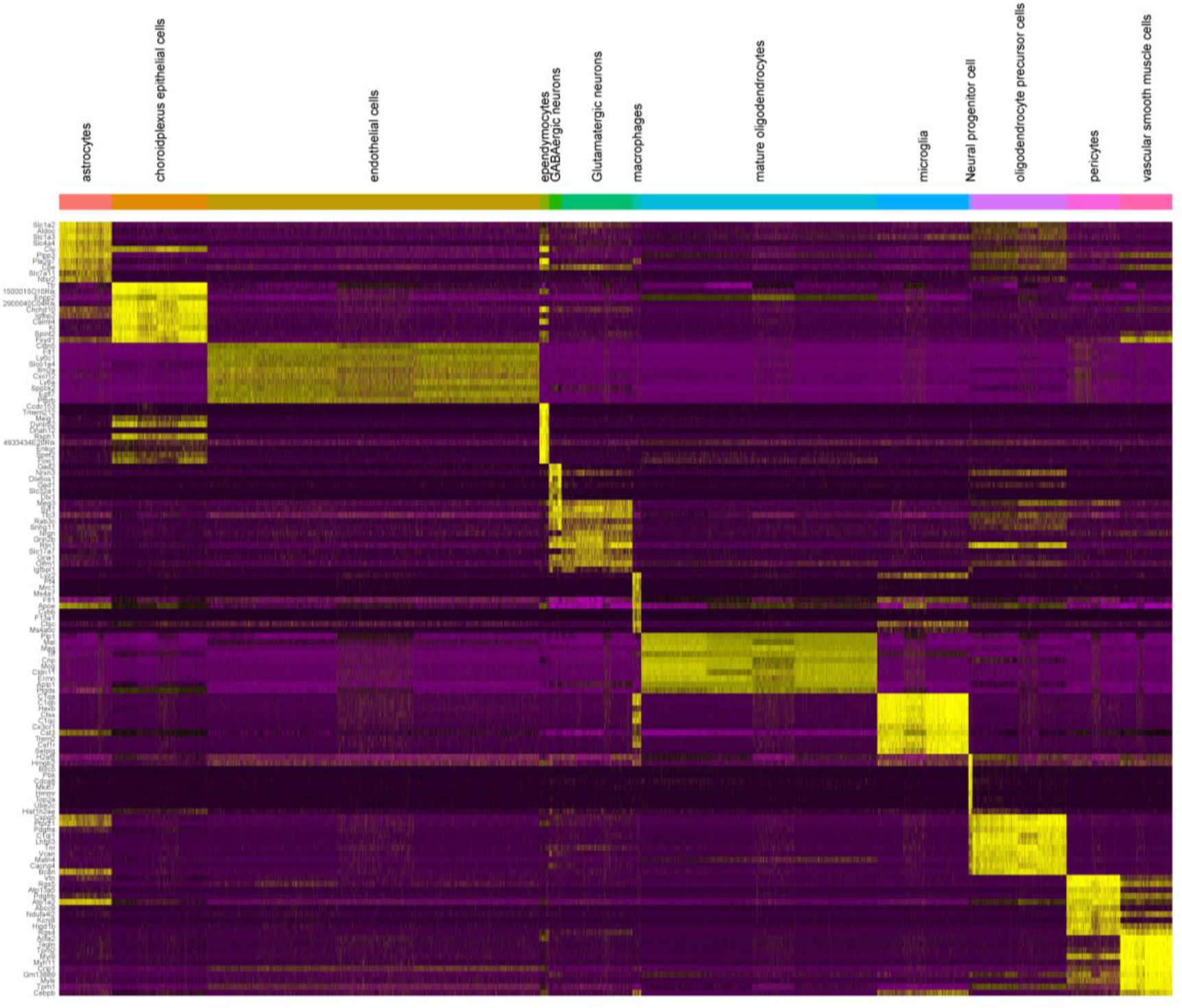
Gene expression heatmap. Heatmap of all 27,069 cells showing the expression levels of the 10 most discriminative genes per cell type (in rows) across all the identified cell populations (in columns). Color-code layout: scale of pink to yellow; from no expression to highest expression.

**Supplementary Fig. 3.**
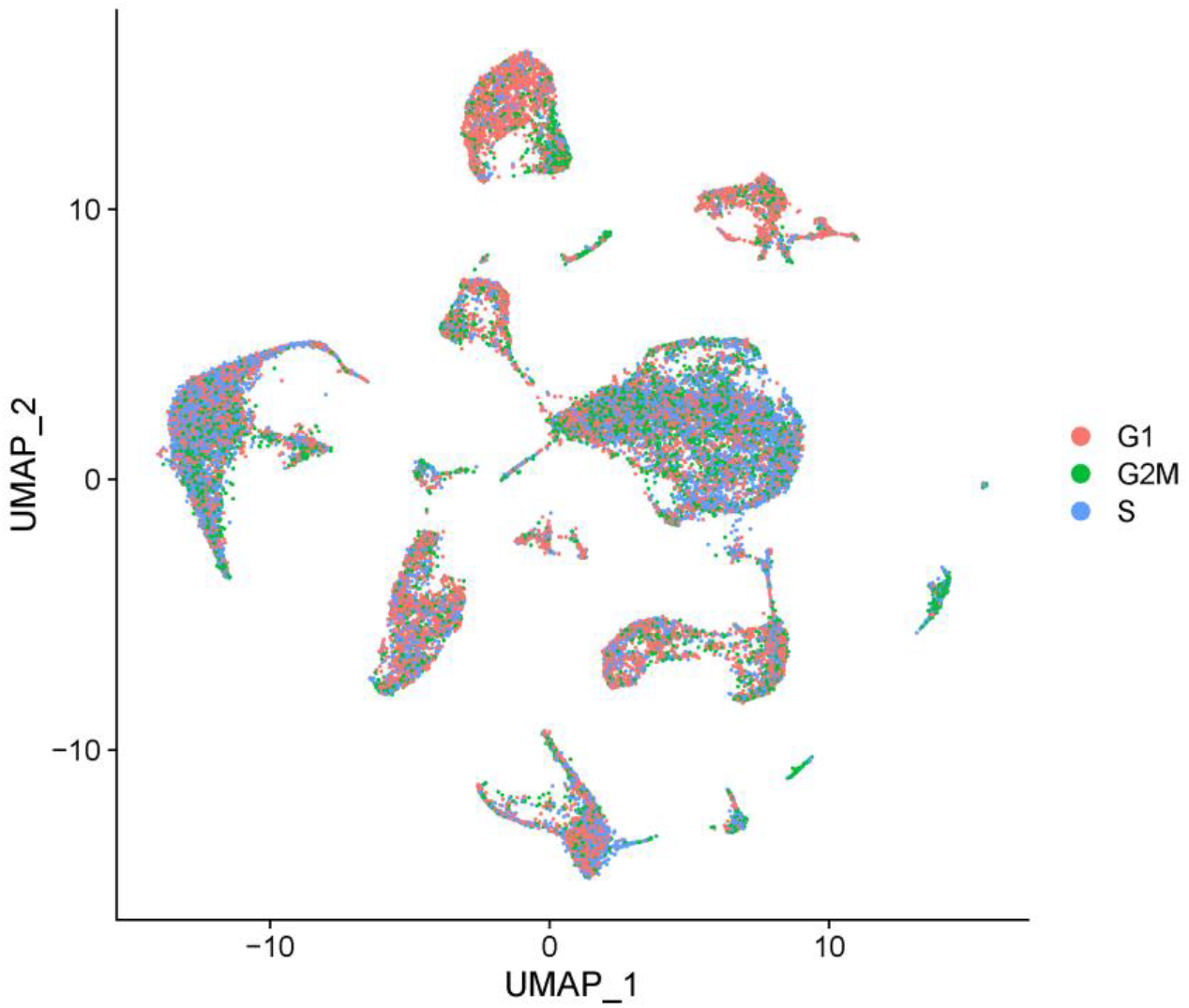
UMAP visualization of cell cycle state in each cell type.

**Supplementary Fig. 4.**
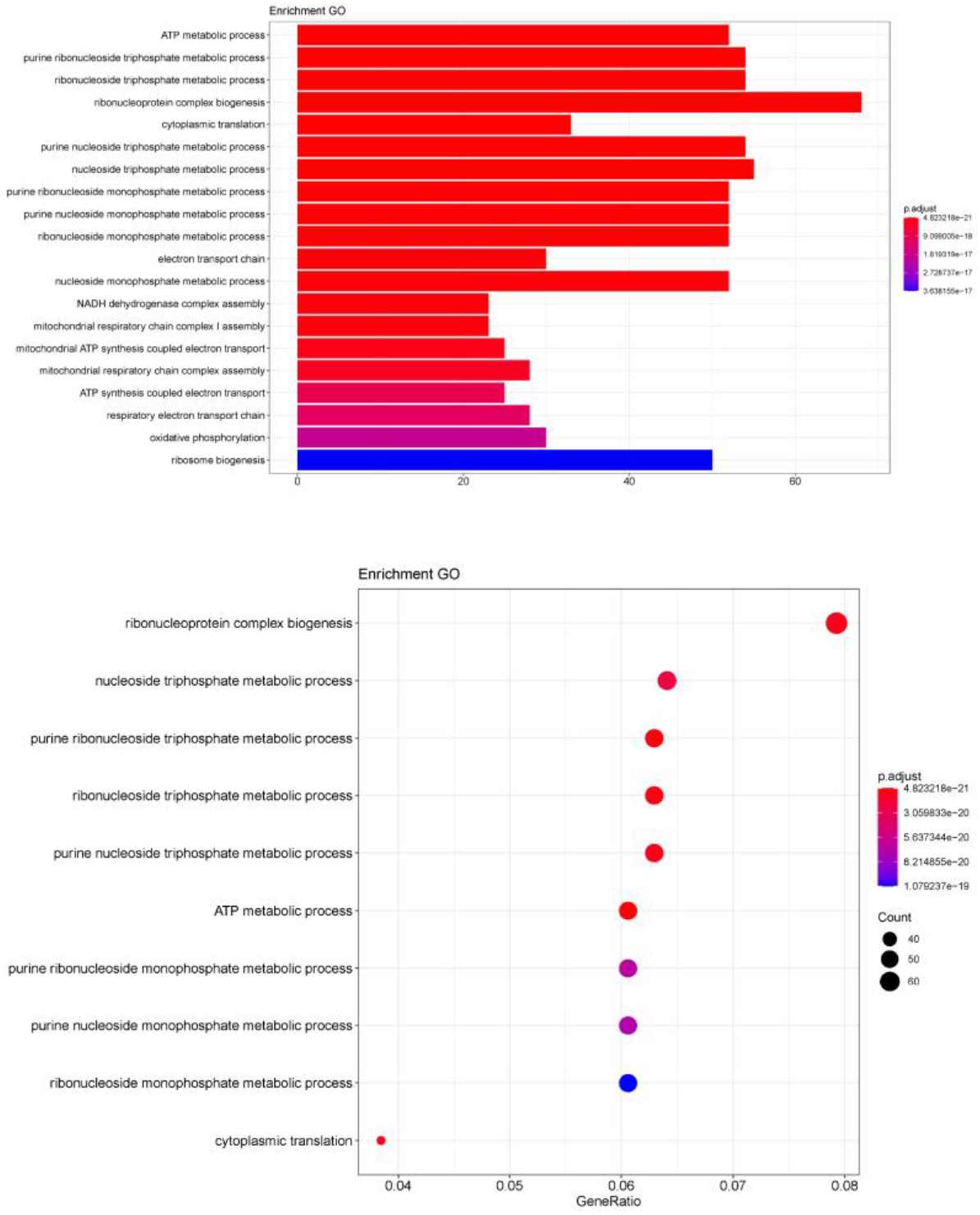
Gene ontology enrichment analysis results in biological process of OPC-2

Supplementary Table 1

Correspondence between cell type and cluster

Supplementary Table 2

List of most discriminating genes per cell type. These cell-type marker genes were calculated by aggregating all sham and PSCI cells identified as belonging to a certain cell type and comparing them to all the other cells. Each cell type has its own worksheet tab within the file with data showing thus: gene name (Gene); the fold change expression (log2FC); original P value (p_val); FDR-adjusted P value (p_val_adj); the fraction of cells transcribing the corresponding gene in the cell type of interest (pct.1); and the fraction of cells transcribing the gene in all the remaining cell types (pct.2).

Supplementary Table 3

Cell type specific DEGs in in ischemic stroke

Supplementary Table 4

Common DEGs in at least 2 cell types

Supplementary Table 5

Numbers of cells in different cell cycle phase of the 13 cell types

Supplementary Table 6

List of most discriminating genes per subcluster of OPC. These subcluster marker genes were calculated by comparing the sham and PSCI cells identified as belonging to a certain subcluster. Each subcluster has its own worksheet tab within the file with data showing thus: gene name (Gene); the fold change expression (log2FC); original P value (p_val); FDR-adjusted P value (p_val_adj); the fraction of cells transcribing the corresponding gene in the subcluster of interest (pct.1); and the fraction of cells transcribing the gene in all the remaining three subclusters (pct.2).

Supplementary Table 7

List of most discriminating genes per cluster of EC. These cluster marker genes were calculated by aggregating all sham and PSCI cells identified as belonging to a certain cluster and comparing them to all the other clusters. Each cluster has its own worksheet tab within the file with data showing thus: gene name (Gene); the fold change expression (log2FC); original P value (p_val); FDR-adjusted P value (p_val_adj); the fraction of cells transcribing the corresponding gene in the cluster of interest (pct.1); and the fraction of cells transcribing the gene in all the remaining clusters (pct.2).

